# Genome-wide screening of circadian and non-circadian impact of Neat1 genetic deletion

**DOI:** 10.1101/2021.01.06.425556

**Authors:** Audrey Jacq, Denis Becquet, Maria-Montserrat Bello-Goutierrez, Bénédicte Boyer, Séverine Guillen, Jean-Louis Franc, Anne-Marie François-Bellan

## Abstract

The functions of the long non-coding RNA, Nuclear enriched abundant transcript 1 (Neat1), are poorly understood. Neat1 is required for the formation of paraspeckles, but its respective paraspeckle-dependent or independent functions are unknown. Several studies including ours reported that Neat1 is involved in the regulation of circadian rhythms. We characterized the impact of Neat1 genetic deletion in a rat pituitary cell line. The mRNAs whose circadian expression pattern or expression level is regulated by Neat1 were identified after high-throughput RNA sequencing of the circadian transcriptome of wild-type cells compared to cells in which Neat1 was deleted by CRISPR/Cas9. The numerous RNAs affected by Neat1 deletion were found to be circadian or non-circadian, targets or non-targets of paraspeckles, and to be associated with many key biological processes showing that Neat1, interacting or independently of the circadian system, could play crucial roles in key physiological functions through diverse mechanisms.

## Introduction

Whereas Nuclear enriched abundant transcript 1 (Neat1) is one of the most studied long non-coding RNA (lncRNA), its roles in physiology and pathology remain still poorly understood. Initially, its physiological role was considered non-essential due to the observation that Neat1 knock-out mice have adequate health and fertility (Nakagawa et al., 2011). It was later found that Neat1 was required for corpus luteum formation affecting fertility in certain sub-populations, and for mammary gland development and lactation in mice (Nakagawa et al., 2014)(Standaert et al., 2014). Furthermore multiple roles for Neat1 in pathology have been since also described, from cancers (Dong et al., 2018) to neurodegenerative (Riva et al., 2016), cardiac (Zhou et al., 2019), immune and viral (Prinz et al., 2019) diseases.

Neat1 is required for the formation of paraspeckles, which are nuclear substructures found in most cultured cells (Hirose et al., 2019). The Neat1 gene is expressed as two variant isoforms: a short one, Neat1_1 (3.8 kb in length in humans) and a long one, Neat1_2 (22.7 kb). Both of these are transcribed from the same promoter, but have different sites of transcriptional termination (Hirose et al., 2019). The longer Neat1 isoform, Neat1_2, is essential for the assembly of paraspeckles, whereas Neat1_1, albeit also a paraspeckle component, is dispensable for their formation and likely plays paraspeckle-independent roles (Li et al., 2017). At the cellular level, Neat1 has been implicated in various processes, including regulation of transcription and chromatin active state and miRNA biogenesis (for review see (Fox et al., 2017) and has been shown to regulate epigenetic marks on histones (Chakravarty et al., 2014)(Butler et al., 2019). The expression of Neat1 has been shown to increase when cells are submitted to a stress (An et al., 2019)(Adriaens et al., 2016)(Adriaens and Marine, 2017). Indeed stress conditions such as hypoxia (Choudhry et al., 2015), viral infection (Imamura et al., 2014), heat shock (Lellahi et al., 2018), mitochondrial stress (Wang et al., 2018) or proteasome inhibition (Hirose et al., 2014) leads to an increase in Neat1.

Although the respective functions of both isoforms as well as the functions that are paraspeckle-dependent or independent remain unclear, it was recently reported in several studies including ours that Neat1 is involved in the regulation of circadian rhythms (Wu et al., 2019)(Chen et al., 2020)(Kukharsky et al., 2020)(Torres et al., 2016a)(Torres et al., 2017).

The mammalian circadian system is a timing system that allows living organisms to anticipate the daily environmental changes, so that behavior and tissue physiology can be adjusted according to the day/night cycle (Asher and Schibler, 2011). The circadian system controls the rhythmicity of a wide range of processes, from molecular (e.g., transcriptional mechanisms), cellular (e.g., the cell cycle (Hong et al., 2014)) physiological (hormonal secretion (Tonsfeldt and Chappell, 2012), metabolic (Bailey et al., 2014)), up to the behavioral processes (sleep/wake cycle, cognitive functions, etc.). This circadian clock system is hierarchically organized and composed of tissue and cellular clocks. Within the cells, a set of clock genes and their protein products, which are highly conserved among animals, generate cell-autonomous rhythms by forming transcription-translation feedback loops through circadian variation in abundance of the positive (Clock and Bmal1) and the negative (Per1, Per2 and Cry1, Cry2) components of the loops. This molecular clock, in turn, drives the rhythmic expression of numerous genes, which ultimately exert control over every biological, physiological, and behavioral process.

Most cellular circadian rhythms are therefore underpinned by daily rhythms in gene expression. While the core circadian system concentrates on transcriptional control, it has been apparent that substantial regulation of clock-controlled genes (ccg) is achieved after transcription so that post-transcriptional controls are emerging as crucial modulators of circadian genes (Lim and Allada, 2013)(Wang et al., 2013)(Menet et al., 2012)(Koike et al., 2012)(Hurley et al., 2014). Indeed in eukaryotes, approximately 1%-10% of genes are subjected to circadian control directly or indirectly but only ~1/5 of the mRNAs that display rhythmic expression are driven directly by transcription, which suggests that post-transcriptional mechanisms including RNA splicing, polyadenylation, mRNA stability, mRNA cytoplasmic export and RNAs nuclear retention are essential layers for generation of gene expression rhythmicity (Partch et al., 2014)(Menet et al., 2012)(Koike et al., 2012)(Hurley et al., 2014)(Torres et al., 2018a)(Mauvoisin, 2019).

We have shown that paraspeckles play a role in the circadian expression of genes through a post-transcriptional mechanism (Torres et al., 2016a). Indeed, paraspeckles are well known to control gene expression at the post-transcriptional level through the nuclear retention of mRNAs (Chen and Carmichael, 2009)(Chen et al., 2008). We have shown in a rat pituitary cell line, the GH4C1 cells, that the expression of numerous paraspeckle components including Neat1 itself, follows a circadian pattern leading to rhythmic variations in paraspeckle number within the cells and that, thanks to their circadian expression pattern and their functions in mRNA nuclear retention, paraspeckles rhythmically retain RNAs in the nucleus. This rhythmic nuclear retention leads to the rhythmic expression of these paraspeckle-target genes (Torres et al., 2016b)(Torres et al., 2017).

A genome-wide screening of the impact of Neat1 deletion was undertaken to evaluate the role of Neat1 on the circadian and non-circadian gene expression in GH4C1 cells. To this end the mRNAs whose circadian expression pattern or expression level is regulated by Neat1 were identified after high-throughput RNA sequencing of the circadian transcriptome of wild type GH4C1 cells compared to that of GH4C1 cells in which Neat1 was deleted by the CRISPR/Cas9 technology. Using Panther analysis (Mi et al., 2019), the biological processes with which the numerous circadian and non-circadian genes affected by Neat1 deletion were associated, were characterized. Finally, whether these genes were or weren’t included in the list of paraspeckle RNA targets previously established (Torres et al., 2016a) was investigated.

## Material and Methods

### Cell line culture

GH4C1 cells, a rat pituitary somatolactotroph line, were obtained in 2012 from ATCC^®^ (CCL-82.2™, lot number: 58945448, Molsheim, France) with certificate analysis and were confirmed to be free of mycoplasma (MycoAlert, Lonza, Levallois-Perret, France). They were grown in HamF10 medium supplemented with 15% horse serum and 2% fetal calf serum.

### CRISPR genomic engineering

For Neat1 knockout, a fragment of 2663nt was edited using a CRISPR/Cas9-based strategy in the 5’ region common to Neat1_1 and Neat1_2 isoforms (Fig. 1A). Two sgRNAs target sites were designed using a CRISPR design tool available on http://crispr.mit.edu and the pairs of oligonucleotides (Supplementary Table 1) were synthetized by Integrated DNA Technologies (Coralville, Iowa, USA). These oligonucleotides were phosphorylated using T4 PNK, annealed and ligated into the *BbsI* (New England Biolabs, Ipswich, MA, USA) restriction site in plasmid pX330-U6-Chimeric_BB-CBh-hSpCas9 (gift from Feng Zhang; Addgene #42230) (Cong et al., 2013). The two sgRNA expression plasmids were amplified using Subcloning Efficiency™ DH5α™ Competent Cells (Thermo Fisher Scientific, Carlsbad, CA, USA). Midi-preparations of plasmids were made with the NucleoBond Xtra Midi kit (Macherey- Nagel, Hoerdt, France) and sequencing was performed (Genewiz, Leipzig, Germany) to confirm the accuracy of the inserted sequence.

**Figure 1.**
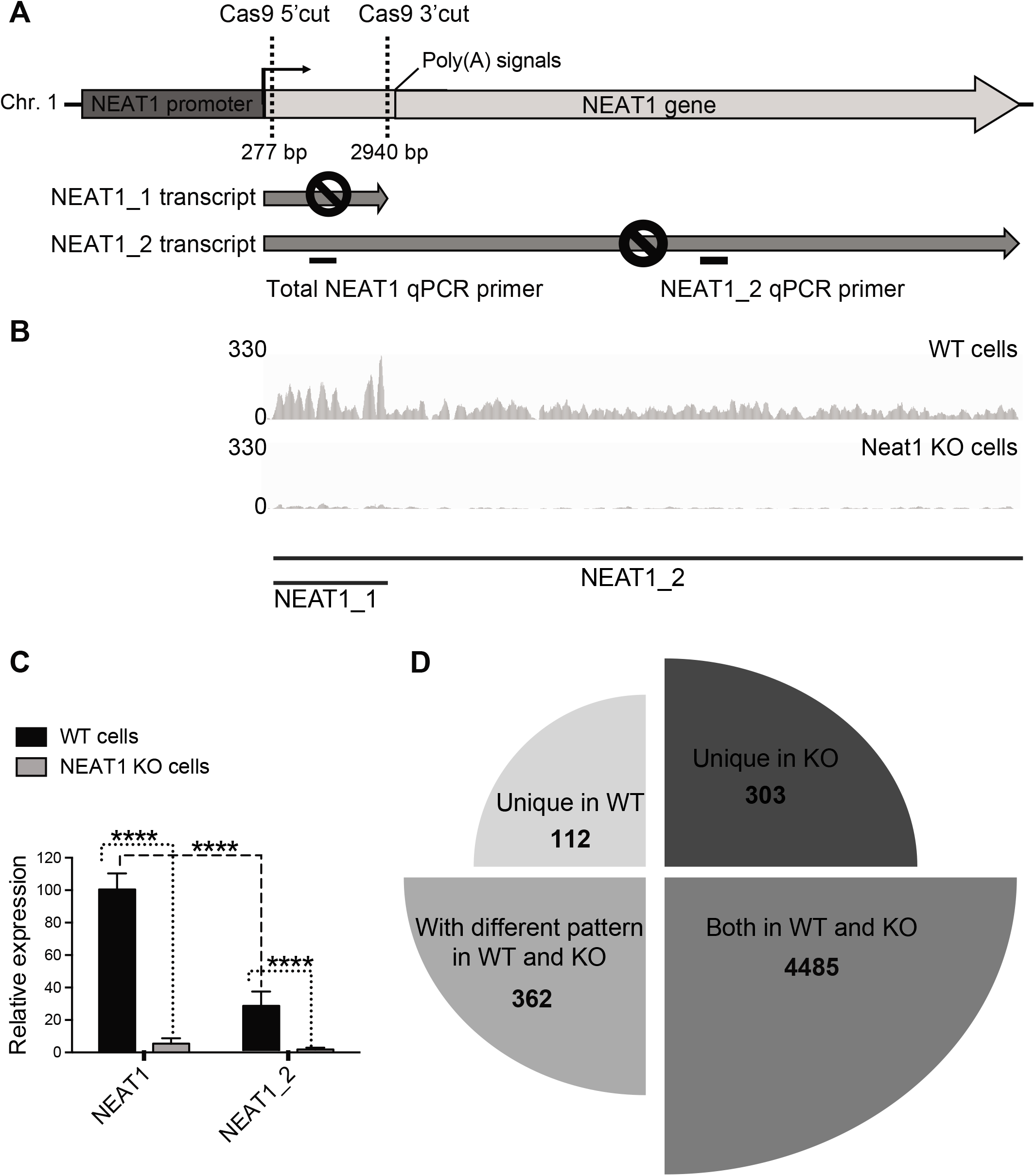
A-C: Schematic representation of Neat1 genetic edition by CRISPR/Cas9 and characterization of Neat1 KO cell line. **A.** The positions of guide RNAs used to delete the Neat1 gene are indicated as well as the positions of qPCR primers used to verify deletion of Neat1 gene (primer sequences are given in Supplementary Table 1). **B**. RNA-Seq peaks for the Neat1 locus that is not officially annotated in the rat genome, demonstrate preferential expression of Neat1_1 in Neat1WT cells and residual Neat1_1 and Neat1_2 expression in Neat1 KO cells. **C.** Significant down-regulation of total Neat1 and Neat1_2 in Neat1 KO GH4C1 lines as revealed by RT-qPCR analysis. qPCR primers (Supplemental Table 1) for total Neat1 were in shared sequences between Neat1_1 and Neat1_2 and detect both isoform RNAs. **D. Impact of Neat1 genetic edition on circadian expressed genes.** Neat genetic deletion leads to a loss of rhythmic pattern in 112 genes (Unique in WT), in a different rhythmic pattern in 362 genes (With different pattern in WT and KO) and in a de novo genesis of rhythmic pattern in 303 genes (Unique in KO). 4485 circadian genes were not affected (Both in WT and KO).

GH4C1 cells were transfected with both plasmids containing sgRNAs, using Lipofectamine 3000 (Thermo Fisher Scientific). Forty-eight hours after the seeding, genomic DNA was extracted using the Nucleospin Tissue kit (Macherey-Nagel) and tested by PCR to screen for genomic editing events, using Hot Start Taq polymerase (New England Biolabs). Subcloning was performed by limit dilution in a 96-well plate. Total RNA of each clonal population was extracted (Macherey-Nagel) and the level of total Neat1 and Neat1_2 expression was monitored by qPCR using specific primers (Supplementary Table 1) whose position was indicated in Fig. 1A. One clone line invalidated for Neat1 (hereinafter referred to as Neat1 KO cells) was selected for the following experiments.

### RNA sequencing experiments

To synchronize cells between themselves, cells were transferred to fresh medium and were harvested after this fresh medium replacement (T0) every 4 h from T10 to T34. For WT and Neat1KO cells, three wells (3.5 cm^2^) were pooled at each time and three replicates were collected from each pool. Total RNA was extracted using the Nucleospin RNA kit (Macherey Nagel). The construction of Illumina DNA libraries and the sequencing from RNA pools were performed by Genewiz (Leipzig, Germany). Libraries were prepared with Illumina Sample Preparation kit with rRNA depletion. Strand-specific RNA-seq was done on Illumina HiSeq 2500, with a read length of 2×150 bp, strand specific (30 million reads per sample on average were obtained).

### RNA-sequencing data processing

Analyses were performed on a local instance of Galaxy. After quality control checks by FastQC and check for adapter content with Trimmomatic, paired-end reads were aligned to the Rat reference genome (Rnor_6.0.80, Ensembl) using STAR (Dobin et al., 2013). The length of the genomic sequence around annotated junctions used was 149 and the other parameters were set to default values. FeatureCounts (Liao et al., 2014) was used to quantify RNA expression at the gene level, using the default values for parameters.

The RNA sequencing data are available at Gene Expression Omnibus (GEO) (accession number n° GSE162751).

### RNA-sequencing analysis

Several tools were used to visually explore sample relationships (Supplemental Figure 1). First, the distribution for logarithm-transformed counts of triplicate of each sample showed a slightly shift for one of the replicates of the T34 time-point in the WT group (Supplemental Figure 1A). However, scatterplot of transformed counts from the two groups at T34 showed that while T34 r1 replicate was less correlated to r2 and r3 in WT than in KO, the correlation was still good enough to keep T34 r1 in the analysis (Supplemental Figure 1B).

### Identification of differentially cycling mRNAs

Rhythmic expression of RNAs was analyzed using ECHO (Extended Circadian Harmonic Oscillators). ECHO is an R powered application to find and visualize circadian rhythms using extended harmonic (De Los Santos et al., 2019). We specified a period of 24h. Differences in rhythmicity between WT and Neat1 KO cells were analyzed using the LimoRhyde (**Li**near **mo**dels for **Rhy**thmicity **de**sign) workflow (Singer and Hughey, 2018). We built a global list of rhythmic genes, applying a 0.1 q-value cutoff and we then used Limma (Ritchie et al., 2015) to characterize the changes of rhythmicity between groups. The q-value cutoff for differentially rhythmic genes was 0.1.

### Identification of differentially expressed mRNAs

The non-rhythmic genes where further analyzed for differential expression using Limma with a 0.01 q-value cutoff.

### Real-Time qPCR

To synchronize cells between themselves, cells were transferred to fresh medium and were harvested after this fresh medium replacement (T0) every 4 h from T2 to T30.

Real time reverse transcriptase PCR (RT-qPCR) was performed using gene-specific primers (Supplemental Table1). The PCR was performed on a CFX96™ Real-Time PCR system using iTaq^TM^ universal SYBR^®^ Green supermix (Bio-Rad, Hercules, CA, USA). Rplp0 and Gapdh were used as endogenous controls. The non-circadian expression for each of those two genes was checked and Rplp0 being the most stable was used for normalizing the expression data of the target genes.

### Cosinor analysis

Cosinor analyses were performed using Prism4 software (GraphPad Software, Inc.). Mean experimental values (± SEM) were fitted using Prism4 by a non-linear sine wave equation: Y = Baseline + Amplitude * sin (Frequency*X + Phase-shift), where Frequency= 2pi/period and period=24h. Goodness-of-fit was quantified using R squared, experimental values being considered well fitted by cosinor regression when the R squared was higher than 0.50. A statistically significant circadian oscillation was considered if the 95% confidence interval for the amplitude did not include the zero value (zero-amplitude test) (Yang et al., 2009) (Kavčič et al., 2011).

## Results

### Loss of Neat1 affects circadian genes in three different ways

Part of the genome corresponding to the Neat1_1 isoform was edited using CRISPR/Cas9 technology (Fig. 1A). RNA-Seq peaks for the Neat1 locus which is not officially annotated in the rat genome, exhibited in WT cells more counts for Neat1_1 compared to Neat1_2 and a residual number of counts for both Neat1_1 and Neat1_2 in Neat1 KO cells (Fig. 1B). Results obtained by RT-qPCR analysis were consistent with RNA-Seq data (Fig. 1C). Indeed the efficacy of CRISPR/Cas9 edition was evaluated by RT-qPCR with primers that detected total Neat1 (Neat1), namely both Neat1_1 and Neat1_2 or only Neat1_2 (Fig. 1A). In Neat1 KO cells, Neat1 as well as Neat1_2 levels were shown to be less than 8% compared to the levels in WT cells (Fig. 1C). Furthermore in WT cells, Neat1_2 was shown to represent around 30% of Neat1 (Fig. 1C). Then, the deletion of the part of the genome of Neat1 corresponding to the Neat1_1 isoform (Fig. 1A) leads also to the decrease in the Neat1_2 expression. This was in agreement with results obtained previously by Yamazaki et al. (Yamazaki et al., 2018).

A global analysis by ECHO, Lymorhyde and Limma applications of the RNA-seq data obtained from both cellular genotypes showed that 5262 genes followed a circadian expression pattern which represented 38% of the 13776 genes expressed in GH4C1 cells (Supplemental Table 2). Since Neat1 is not officially annotated in the rat genome, it was not included in the list of genes that displayed a rhythmic pattern, but we showed by an additional qPCR analysis that Neat1 as well as Neat1_2 exhibited a circadian expression pattern in WT cells, as we previously reported (Supplemental Figure 2). This was not only consistent with our previous report (Torres et al., 2016a), but this corroborated another study that has also reported since there the circadian expression of Neat1 in skin fibroblasts (Ray et al., 2020).

Among the 5262 genes with a circadian expression pattern, 777 (14.8%) exhibited circadian differences between the two cellular genotypes (Fig. 1D; Supplemental Table 2). It was observed that 112 genes (2%) became arrhythmic when Neat1 was deleted, 362 genes (6.9%) displayed a modified circadian pattern and 303 genes (5.7%) acquired a circadian pattern after Neat1 editing (Fig. 1D and Fig. 2; Supplemental Table 3). It then appeared that while around 15% of rhythmic genes in GH4C1 cells displayed a circadian pattern dependent on Neat1, these Neat1-dependent circadian genes could be classified in three categories depending on whether they lost their circadian pattern, they displayed a modified one or they acquired one (Fig. 1D and Fig. 2).

**Figure 2.**
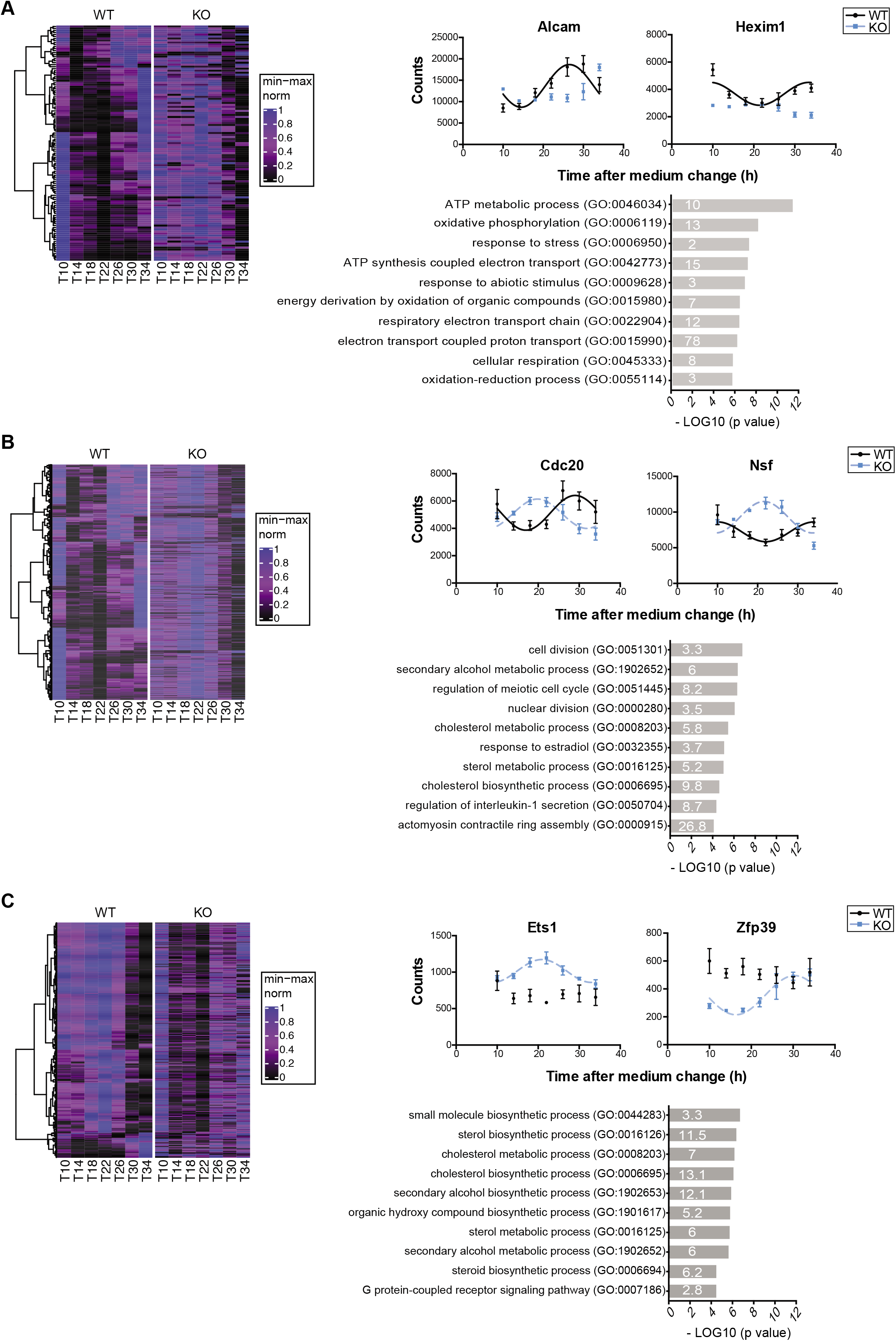
Analysis of circadian genes affected by Neat1 deletion. A. Analysis of genes from the category “Unique in WT”. **Left Panel** Heat map representation of genes found to belong to the category “Unique in WT”. Heat maps have been built using the euclidean distance as a measurement of how similar gene expressions in WT group are to each other and the ward.D2 method to group them based on that distance. Corresponding genes are displayed in the same order in KO group. The dendrogram obtained from hierarchical clustering is shown on the left side of the heatmap. A min-max normalization was applied for each gene independently and is represented by colors on the heatmap: higher, blue; lower, black. The hour of cell collection is indicated below (T). **Right Upper Panel** Cosinor analysis of RNA- Seq counts of two genes, Alcam and Hexim1, shows that counts from WT cells could be adequately fitted (R^2^>0.50) with a non-linear cosinor equation in which the period value was set to 24 h (cosinor fit values given in Supplemental Table 4), while counts from Neat1 KO cells can’t. At each time point, data are means ± SEM of samples from the three biological replicates of RNA-Seq. **Right Lower Panel** Functional characterization by Panther Analysis showing the Gene Ontology biological processes with the highest p value. The numbers inside the columns correspond to the fold enrichment. **B. Analysis of genes from the category “With different pattern in WT and KO”. Left Panel** Heat map representation of genes found to belong to the category “With different pattern in WT and KO”. Heat maps have been built using the euclidean distance as a measurement of how similar gene expressions in WT group are to each other and the ward.D2 method to group them based on that distance. Corresponding genes are displayed in the same order in KO group. The dendrogram obtained from hierarchical clustering is shown on the left side of the heatmap. A min-max normalization was applied for each gene independently and is represented by colors on the heatmap: higher, blue; lower, black. The hour of cell collection is indicated below (T). **Right Upper Panel** Cosinor analysis of RNA-Seq counts of two genes, Cdc20 and Nsf, shows that counts from WT and KO cells could be adequately fitted (R^2^>0.50) with a non-linear cosinor equation in which the period value was set to 24 h (cosinor fit values given in Supplemental Table 4). At each time point, data are means ± SEM of samples from the three biological replicates of RNA-Seq. **Right Lower Panel** Functional characterization by Panther Analysis showing the Gene Ontology biological processes with the highest p value. The numbers inside the columns correspond to the fold enrichment. **C. Analysis of genes from the category “Unique in KO”. Left Panel** Heat map representation of genes found to belong to the category “Unique in KO”. Heat maps have been built using the euclidean distance as a measurement of how similar gene expressions in KO group are to each other and the ward.D2 method to group them based on that distance. Corresponding genes are displayed in the same order in WT group. The dendrogram obtained from hierarchical clustering is shown on the left side of the heatmap. A min-max normalization was applied for each gene independently and is represented by colors on the heatmap: higher, blue; lower, black. The hour of cell collection is indicated below (T). **Right Upper Panel** Cosinor analysis of RNA-Seq counts of two genes, Ets1 and Zfp39, shows that counts from KO cells could be adequately fitted (R^2^>0.50) with a non-linear cosinor equation in which the period value was set to 24 h (cosinor fit values given in Supplemental Table 4), while counts from WT cells can’t. At each time point, data are means ± SEM of samples from the three biological replicates of RNA-Seq. **Right Lower Panel** Functional characterization by Panther Analysis showing the Gene Ontology biological processes with the highest p value. The numbers inside the columns correspond to the fold enrichment.

We previously showed that another lncRNA, Malat1, which is not officially annotated in the rat genome is closely associated with Neat1 (Torres et al., 2018b). Malat1 was shown here in an additional qPCR analysis to belong to the first category of Neat1 regulated genes since Malat1 exhibited a circadian expression pattern in WT cells that was lost after Neat1 deletion (Supplemental Figure 2). This altered circadian pattern of Malat1 was further associated with a decrease in its expression level (Supplemental Figure 2). Since it has been shown that the expression of Malat1 is not altered in different tissues of mice lacking the Neat1 poly-adenylation signal necessary for the formation of the short isoform of Neat1 (Isobe et al., 2020), it may be assumed that the lack of Neat1_2 was specifically responsible for the effects on Malat1 we reported here.

Within two of the three categories of Neat1 gene regulation mentioned above, we found some core-clock genes. This was the case for Per2 whose circadian expression pattern was lost in Neat1 KO cells (Supplemental Figure 3) and for Per1, Per 3 and Cry 1 whose circadian expression pattern differed between the two cell lines (Supplemental Figure 3). Moreover direct clock-controlled genes considered as transcriptional relays between the molecular core-oscillator and target genes such as Tef or Nfil3 were also shown to display a different circadian pattern in Neat1 KO cells as compared to WT cells (Supplemental Figure 3).

Within each of the three categories of genes, we isolated 3 to 4 clusters at the third node (Fig. 2A-C Left Panels). Whereas oscillating patterns were consistent within each cluster, they were distinct between clusters, with transcripts peaking at different times during the 24h cycle (Fig. 2A-C Left Panels). Several genes from each of these three categories were selected and their circadian pattern fitted by cosinor analysis using either RNA-seq counts (Fig. 2A-C Right Upper Panels) or qPCR measurements (Supplemental Figure 4). Accordingly two genes (Alcam and Hexim1) were shown here to lose their circadian expression pattern in Neat1 KO cells. Indeed while counts from WT cells could be adequately fitted (R^2^>0.50) with a nonlinear cosinor equation in which the period value was set to 24 h, counts from Neat1 KO cells couldn’t (Fig. 2A Right Upper Panel; cosinor fit values given in Supplemental Table 4). The same hold true for two others genes selected from the same category (Nos1 and Mtnd4) whose expression pattern was determined after RT-qPCR analysis (Supplemental Figure 4). While being adequately fitted (R^2^ >0.50) with a non-linear cosinor equation with a period value of 24 h in both WT and Neat1 KO cells, the genes Cdc20 and Nsf (Fig. 2B Right Upper Panel; cosinor fit values given in Supplemental Table 4) as well as the genes Aura and Ccnb1 (Supplemental Figure 4) displayed a phase inverted pattern between the two genotypes. As examples of genes belonging to the third category, we showed that Ets1 and Zfp39 were rhythmic in Neat1 KO and not in WT cells (Fig. 2C Right Upper Panel; cosinor fit values given in Supplemental Table 4). This was also the case for Mvk and Fdft1 as shown in Supplemental Figure 4. Whether these three categories of circadian genes that were differently affected by Neat1 deletion were involved in different functions was further assessed by Panther analysis (Mi et al., 2019).

### Circadian genes impacted by Neat1 deletion are associated with different functions

The circadian genes becoming arrhythmic after Neat1 deletion were shown to be mainly involved in energy metabolism and response to stress after Panther analysis (Fig. 2A Right Lower Panel, Supplemental Table 5). We indeed found that in the list of genes from the first category, namely genes that displayed a circadian expression pattern only in WT, the terms “energy metabolism” and “response to stress” were among the most prominent enriched annotation clusters in Gene Ontology biological processes (Fig. 2A Right Lower Panel). These functional annotations were consistent with the recently reported cross-regulation between paraspeckles and mitochondria (Wang et al., 2018) and with the numerous studies showing that Neat1 acts as an important sensor and effector during stress (Lellahi et al., 2018) (Adriaens and Marine, 2017).

The circadian genes with a Neat1 dependent cycling profile were further mainly involved in cell cycle and sterol metabolism. Functional classification of genes from the second category, namely genes with a different circadian pattern in Neat1 KO cells showed indeed the prominence of annotations such as “cell division” and “sterol metabolic processes” (Fig. 2B Right Lower Panel, Supplemental Table 5). The genes from this category could then constitute the substratum of the implication of Neat1 in proliferative processes and in lipid metabolism (Wang, 2018) (Sun et al., 2019). Indeed Neat1 has been not only involved in the formation of physiological tissue such as the mammary gland or the corpus luteum (Standaert et al., 2014) (Nakagawa et al., 2014) but its deregulation has been extensively described in tumor processes (for a review Yu et al., 2017). Finally circadian genes acquiring a circadian pattern after Neat1 deletion were also mainly involved in “sterol metabolic processes” and in “G protein-coupled receptor signaling pathway” (Fig. 2C Right Lower Panel, Supplemental Table 5).

### Down-regulated and up-regulated genes after Neat1 deletion are functionally distinct

Apart from circadian expressed genes, 3262 genes were found differentially expressed following Neat1 deletion representing near 25% of genes expressed in GH4C1 cells (13376 genes). Among these 3262 genes, 1354 were up- and 1908 down-regulated (Fig. 3A, 3B; Supplemental Table 2). Panther analysis of the top 1,000 up- or down-regulated genes was carried out. This analysis revealed distinct gene classes whose expression was altered by Neat1 deletion. Down-regulated genes were shown to be mainly involved in “immune processes” (Fig. 3C, Supplemental Table 6) consistent with the well recognized view that Neat1 is an immunity-associated lncRNA that regulates important cytokine production and the immune response (Zhang et al., 2016a) (Ye et al., 2020). Up-regulated genes were themselves merely associated with “regulation of gene expression” and “RNA biosynthetic process” (Fig. 3E, Supplemental Table 6). A few genes from both lists of up- and down- regulated genes were selected and their RNA-seq counts reported during the circadian period to illustrate down-regulated genes associated with “immune processes” and up-regulated genes associated with “regulation of gene expression”, (Fig. 3D, 3F).

**Figure 3.**
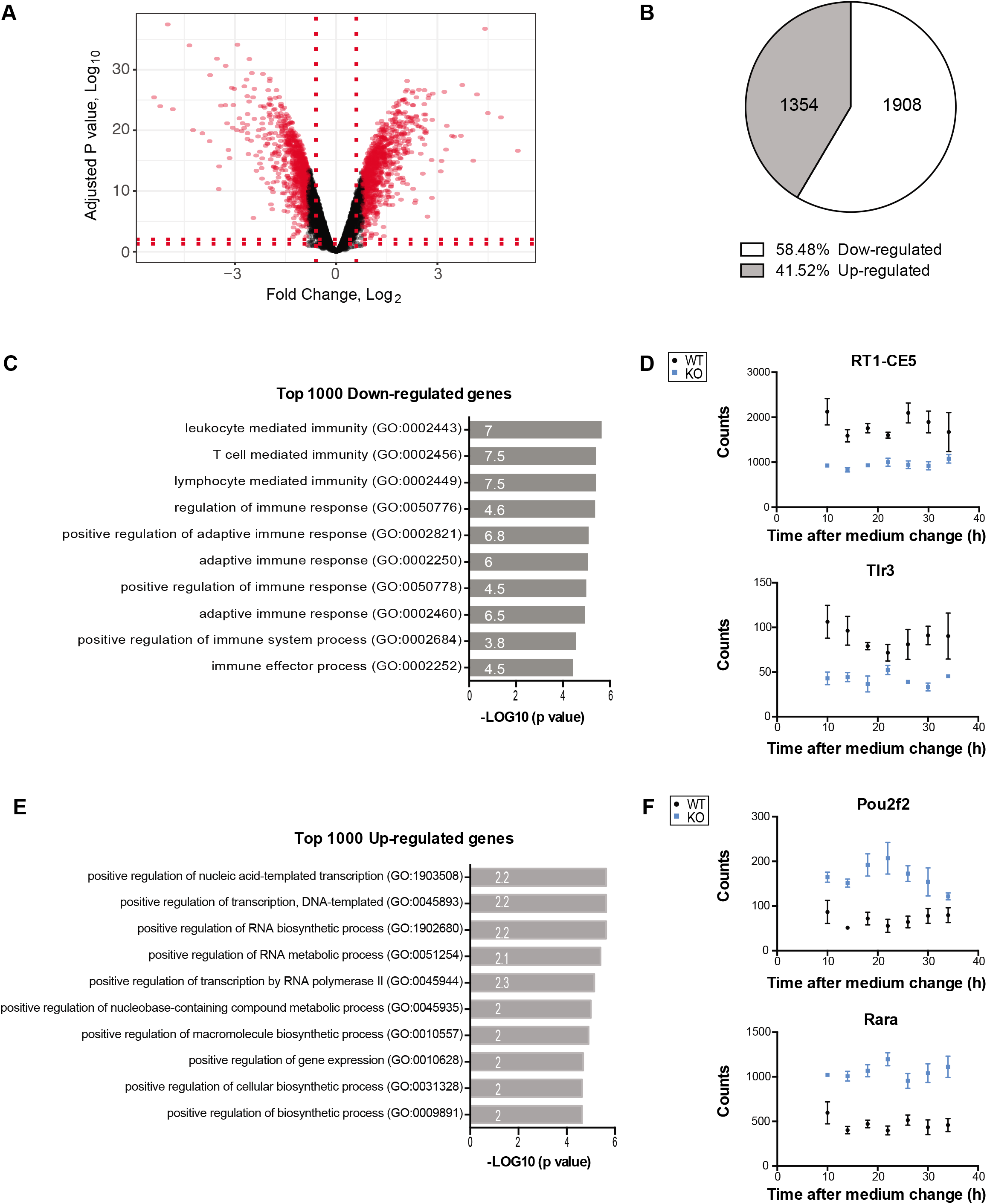
Analysis of down-regulated and up-regulated genes after Neat1 deletion. **A.** Volcano plot for Neat1 KO cells versus WT cells is shown; the top 1000 down-regulated and the top 1000 up-regulated genes are highlighted in red. **B**. The proportion of down-regulated and up-regulated genes is also shown (gene numbers are given in pie charts). **C.** Functional characterization by Panther Analysis of the top 1000 down-regulated genes showing the Gene Ontology biological processes with the highest p value. The numbers inside the columns correspond to the fold enrichment. **D.** RNA-Seq counts of two genes, RT1-CE5 and Tlr3, from the top 1000 down-regulated genes are given as an example. At each time point, data are means ± SEM of samples from the three biological replicates of RNA-Seq. **E.** Functional characterization by Panther Analysis of the top 1000 up-regulated genes showing the Gene Ontology biological processes with the highest p value. The numbers inside the columns correspond to the fold enrichment. **F.** RNA-Seq counts of two genes, Pou2f2 and Rara, from the top 1000 up-regulated genes are given as an example. At each time point, data are means ± SEM of samples from the three biological replicates of RNA-Seq.

### A quarter of circadian expressed genes are paraspeckle RNA targets

We previously determined the list of RNAs that are paraspeckle targets (Torres et al., 2016a). Since the circadian variation of the number of paraspeckles within the nucleus of GH4C1 cells we reported (Torres et al., 2016a) is believed to account for the circadian expression of paraspeckle RNA targets, we compared the list of paraspeckle RNA targets, we previously established, with that of genes that displayed a circadian expression pattern presented here. We found that among the 5262 genes that displayed a circadian expression pattern, 1380 (26%) were paraspeckle targets (Fig. 4A). Since a quarter of circadian expressed genes were associated with paraspeckles, the contribution of the paraspeckles to the circadian transcriptome was believed very significant.

**Figure 4.**
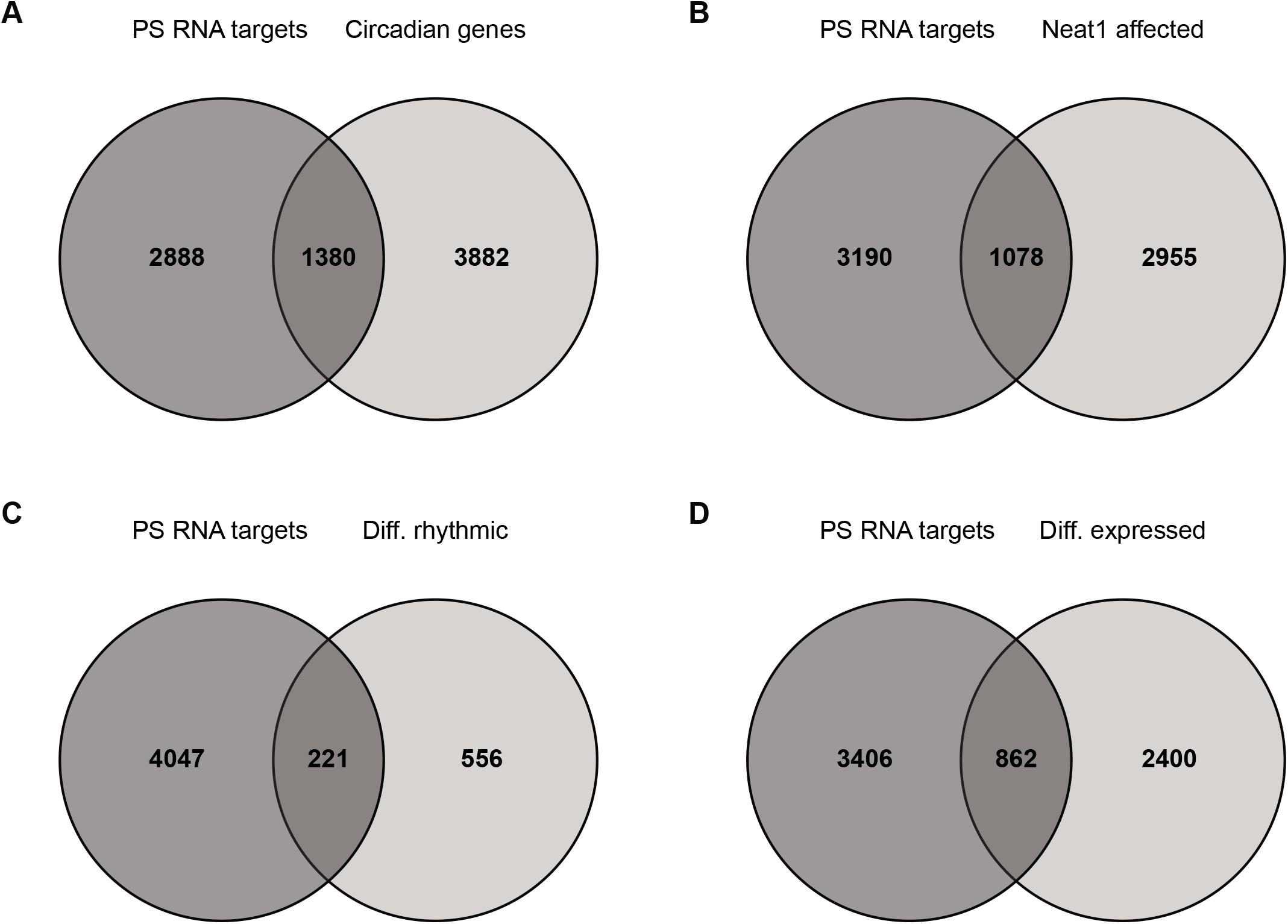
A. Contribution of Paraspeckle RNA targets to the circadian transcriptome. Venn diagram representing overlaps between circadian transcripts and Paraspeckle (PS) RNA targets. **B-D. Contribution of Paraspeckle RNA targets to Neat1 dependent transcriptome.** Venn diagrams representing overlaps between Paraspeckle (PS) RNA targets and total genes affected by Neat1 deletion (B), circadian genes affected by Neat1 deletion (C) and genes whose expression is affected by Neat1 deletion (D).

### A quarter of genes affected by Neat1 deletion are paraspeckle RNA targets

When taking into account both the genes whose expression level was modified by Neat1 deletion (3262 genes) and those whose circadian expression was affected (777 genes), it was concluded that Neat1 was an important regulator since near 30% of genes expressed by GH4C1 cells (13776 genes) were affected by Neat1 deletion.

Since Neat1 deletion is believed to disrupt paraspeckle bodies, we evaluated the number of genes affected by Neat1 deletion (i.e. whose circadian expression or expression level was modified) with that of genes associated with paraspeckles, by overlapping this list with that of paraspeckle RNA targets, we previously reported (Torres et al., 2016a). Neat1 deletion affected 1078 genes (26%) that were paraspeckle RNA targets (Fig. 4B). We further discriminated among these 1078 genes those that displayed a modified circadian pattern from those that exhibited an altered expression level. We found that among the 777 genes that displayed an altered circadian pattern after Neat1 deletion, 221 (28%) were paraspeckle RNA targets (Fig. 4C) and among the 3262 genes with an altered expression level after Neat1 deletion, 862 genes (26%) were paraspeckle RNA targets (Fig. 4D). It then appeared that the proportion of genes affected by Neat1 deletion that were associated with paraspeckles, was equivalent whether Neat1 regulated their level of expression or their circadian expression.

## Discussion

The present study shows that in GH4C1 cells, the circadian transcriptome turns out to be really significant representing no less than 38% of expressed genes. Of course the use of ECHO to identify rhythmic genes, an application that has been shown to out-perform existing approaches to identify oscillations in genome-scale circadian datasets (De Los Santos et al., 2019) may explain the great proportion of rhythmically expressed genes found here. Indeed, the development of high-throughput sequencing techniques (Hughes et al., 2012) coupled to always more efficient and robust methods to detect rhythmic expression profiles has led to the re-evaluation of the extent of the circadian transcriptome in a given tissue or cell line. Depending on the tissue or cell type studied, the proportion of rhythmically expressed genes is estimated to span from 6 to 20%. Moreover when considering the whole organism, it was first estimated that approximately half of the genome is oscillating somewhere in the body (Zhang et al., 2014), but nowadays substantial improvements in performance have amplified the notion that circadian oscillations are pervasiveness by showing that 93.5% of all possible protein coding transcripts exhibit circadian oscillations in at least one tissue (Ceglia et al., 2018). In any case, the significant extent of the circadian transcriptome in GH4C1 cells is not only consistent with our previous reports showing that these cells harbor a functional circadian oscillator (Guillaumond et al., 2011) (Torres et al., 2016a) but also supports the relevance to use this cell line for circadian issues.

Although there are various computational methods to detect rhythmicity in genome-scale data, methods to detect changes in rhythmicity or changes in average expression between different conditions are scarcer. We used LimoRhyde a method that allows detecting changes in rhythmicity across different conditions and therefore allows unraveling the effects of genetic perturbations on biological rhythms (Singer and Hughey, 2018). In keeping with the view that Neat1 is deeply involved in the circadian system functioning (Wu et al., 2019) (Chen et al., 2020) (Kukharsky et al., 2020) (Torres et al., 2016a) (Torres et al., 2017), this computational method allows us to show that the CRISPR/Cas9 editing of Neat1 in GH4C1 cells affects no less than 15% of circadian expressed genes. While the number of genes that became arrhythmic following Neat1 deletion is relatively weak, a more significant number of genes display a modified circadian profile in Neat1 KO cells. The same hold also true for core-clock genes themselves. Whereas only Per2 loses its circadian expression pattern, several other core-clock genes as well as their direct targets such as transcription factors known to relay circadian oscillations to circadian-controlled genes, display a modified circadian profile after Neat1 removal. Of course these core-clock genes and their direct targets that are altered after Neat1 removal may trigger part of the observed modifications in circadian-clock-controlled genes. Strikingly, in addition to disrupting the circadian expression of numerous genes that become either arrhythmic or display a different profile, Neat1 deletion causes an unexpected genesis of de novo oscillating transcripts. Whether these de novo oscillating transcripts are genes whose circadian expression in WT cells is masked by an anti-phasic circadian rhythm generated by Neat1 or whether they reflect a real reprogramming of the circadian transcriptome after Neat1 deletion remains to be determined. Whatever the case, when analyzed for singular enrichment in biological processes, the circadian genes affected by Neat1 deletion exhibited different functional association depending on whether they lost or gain rhythmicity or whether they display a modified profile.

Interestingly, genes that lose their circadian expression after Neat1 deletion were found associated with both “energy metabolism” and “response to stress”. This is reminiscent of the well recognized involvement of Neat1 in the cellular response to stress (An et al., 2019) (Adriaens et al., 2016) (Adriaens and Marine, 2017) as well as in the cross-regulation that has been reported between paraspeckles and mitochondria (Wang et al., 2018). Indeed it has been shown that Neat1 depletion has important effects on mitochondrial dynamics by altering the sequestration of mitochondrial mRNAs in paraspeckles (Wang et al., 2018). It should be noted that the interaction between the circadian clock and the response to stress is well established and that the interest in regulation of mitochondria by the circadian timekeeping system has gained interest as more and more evidence indicates that the biological clock orchestrates the functioning of mitochondria and suggests that the circadian clock coordinates energy metabolism and cellular anti-oxidant mechanisms that prevent oxidative damage (de Goede et al., 2018) (Sardon Puig et al., 2018). In view of the present findings it is then tempting to speculate that Neat1 could be one of the links through which the circadian clock is involved in both the response to stress and the regulation of energy metabolism.

The numerous genes that exhibit a modified circadian pattern after Neat1 deletion were rather shown associated with “cell cycle” and “sterol metabolism”. Interestingly the genetic ablation of Neat1 in mice results in disturbances of cell growth such as aberrant mammary gland morphogenesis (Standaert et al., 2014) or impaired corpus luteum formation associated with a lack of progesterone synthesis (Nakagawa et al., 2014). In addition, Neat1 is now well recognized as a critical component in the progress and development of cancer (Pisani and Baron, 2020). Of note it has now become obvious that cholesterol pathways are intertwined with circadian clock and that the circadian clock affects the carcinogenesis by regulating the lipid metabolism (Kovač et al., 2019). The finding we report here that Neat1 is involved in the regulation of clock-controlled-genes associated both with “cell cycle” and “sterol metabolism” suggests that Neat1 plays a role in the interconnection of the circadian system with cell growth.

While 15% of circadian-expressed-genes are shown here altered in Neat1 KO cells, 25% of non-circadian genes exhibit a modified level of expression. This strengthens the regulatory role exerted by Neat1 on GH4C1 transcriptome. Of note the top down-regulated genes are mostly associated with “immune processes” in keeping with several reports highlighting the emergence of Neat1 as an important regulatory layer in controlling gene expression in innate immunity and adaptive immune response (Zhang et al., 2016b) (Zhang et al., 2019). In particular Neat1 is considered to be critical for the immune response against virus (Prinz et al., 2019). It is therefore not surprising to find here that Neat1 positively regulates numerous genes closely associated with immune functions such as Tlr3 which is one of the Toll□like receptors (TLRs) that are central players in the early host immune response to acute viral infection and have been shown to play a crucial role in defense against microorganisms (Zeng et al., 2019). Consistent with the positive regulation of Tlr3 by Neat1 we report here, Neat1 has been recently found inversely correlated to the level of Tlr3 in patients with Hepatitis B virus infection (Zeng et al., 2019).

Neat1 through its binding to an RNA-binding protein, Hexamethylene bis-acetamide-inducible protein 1 (Hexim1) has been shown to play an important role in regulating DNA-mediated induction of the innate immune response (Morchikh et al., 2017). It is then of note that in GH4C1 cells, Hexim1 follows a circadian expression pattern which is under Neat1 control since this pattern is lost after Neat1 removal. Importantly a number of immune functions follow diurnal variations including for instance, counts of lymphocytes, T lymphocytes, and B lymphocytes in human blood (Cermakian et al., 2013), cytokine and chemokine expression (Rahman et al., 2015), response to antigen presentation (Fortier et al., 2011) or leukocyte tissue recruitment (Scheiermann et al., 2012). Whether the crosstalk between Neat1 and the immune functions implies the rhythmicity of the innate immune response remains to be explored.

The comparison of the paraspeckle RNA targets we previously established in GH4C1 cells (Torres et al., 2016a) with circadian-expressed genes identified here allows us to show that the circadian nuclear retention by paraspeckles of mRNAs contributes significantly to the circadian transcriptome in this cell line. Surprisingly however the proportion of genes that in addition to be affected by Neat1 deletion, are also paraspeckle RNA targets, is the same in the non-circadian gene group compared to the circadian one and represents only a quarter of Neat1 regulated genes. Therefore three inferences may be drawn from these data. First the paraspeckle RNA targets may be arrhythmic, whether they are or they aren’t affected by Neat1 deletion. It may then be that the rhythmic pattern generated by paraspeckle rhythm for those RNAs may be counterbalanced by an opposite rhythm in their transcription. As an alternative, the quantity of these RNAs associated to paraspeckles may not be sufficient to generate their rhythmic pattern. The second inference drawn from our data is that some paraspeckle RNA targets are not affected by Neat1 deletion suggesting that the association of these RNAs with paraspeckles is not dispensable for their expression. Finally since there are both rhythmic and non-rhythmic genes affected by Neat1 that aren’t paraspeckle targets, it may be suggested that some Neat1 regulation exerted on the rhythmic pattern or on the expression level of RNAs may be independent of paraspeckles. In this view, there are several ways by which Neat1 can regulate the expression of genes that aren’t associated with paraspeckles. First acting as a miRNA sponge, Neat1 can mediate gene regulation through paraspeckle independent mechanisms. This mechanism is utilized in various cancers such as hepatocellular carcinoma (Li et al., 2020). Another way, through which Neat1 has been shown to regulate the transcription of some genes, is via the retention of proteins such as transcription factors in paraspeckles. For instance the retention of the transcriptional and splicing regulator SFPQ in paraspeckles impedes the ability of SFPQ to bind the chromatin of target genes, leading to changes in the transcriptional output, such as the repression of Adarb2 (Hirose et al., 2014) and the induction of Il8 (Imamura et al., 2014). Neat1 can also direct gene repression through DNA methylation of gene promoters (Zhang et al., 2018) and has been shown to alter chromatin marks such as histone H3K4 trimethylation and histone H3K9 acetylation in specific genes (Chakravarty et al., 2014). Neat1 could play a role in the control of chromatin remodeling. Proteins from the SWI/SNF chromatin remodeling complex were shown to be major components of the paraspeckles (Kawaguchi et al., 2015) leading to the assumption that Neat1 could alter chromatin organization by bringing these proteins to the chromatin of active genes or by retaining them in the paraspeckles. Finally, as both isoforms of Neat1 have been deleted in our KO cells, the Neat1 regulation of genes that are not associated with paraspeckles may be attributed to the deletion of the short isoform that is known to have paraspeckle independent roles (Li et al., 2017) (Naveed et al., 2020).

In summary, the genetic deletion of Neat1 in GH4C1 cells affects many circadian- and non-circadian genes that, depending on the type of control exerted by Neat1, are associated with different specific biological processes, including “energy metabolism”, “response to stress”, “cell division”, “sterol metabolism”, “immune processes” and “gene regulation”. Furthermore, by overlapping the list of genes affected by Neat1 deletion with that of paraspeckle RNA targets previously established in this cell line (Torres et al., 2016a), the existence of different mechanisms through which Neat1 can exerts its regulatory control is proposed. Overall, the present data shed light on the crucial roles Neat1 can play through multiple diverse mechanisms in cellular biological processes.

## Notes

https://www.ncbi.nlm.nih.gov/geo/query/acc.cgi?acc=GSE162751

### Competing Interest Statement

The authors have declared no competing interest.

