## Supplemental Figure 1 for "Genome-wide screening of circadian and non-circadian impact of Neat1 genetic deletion"

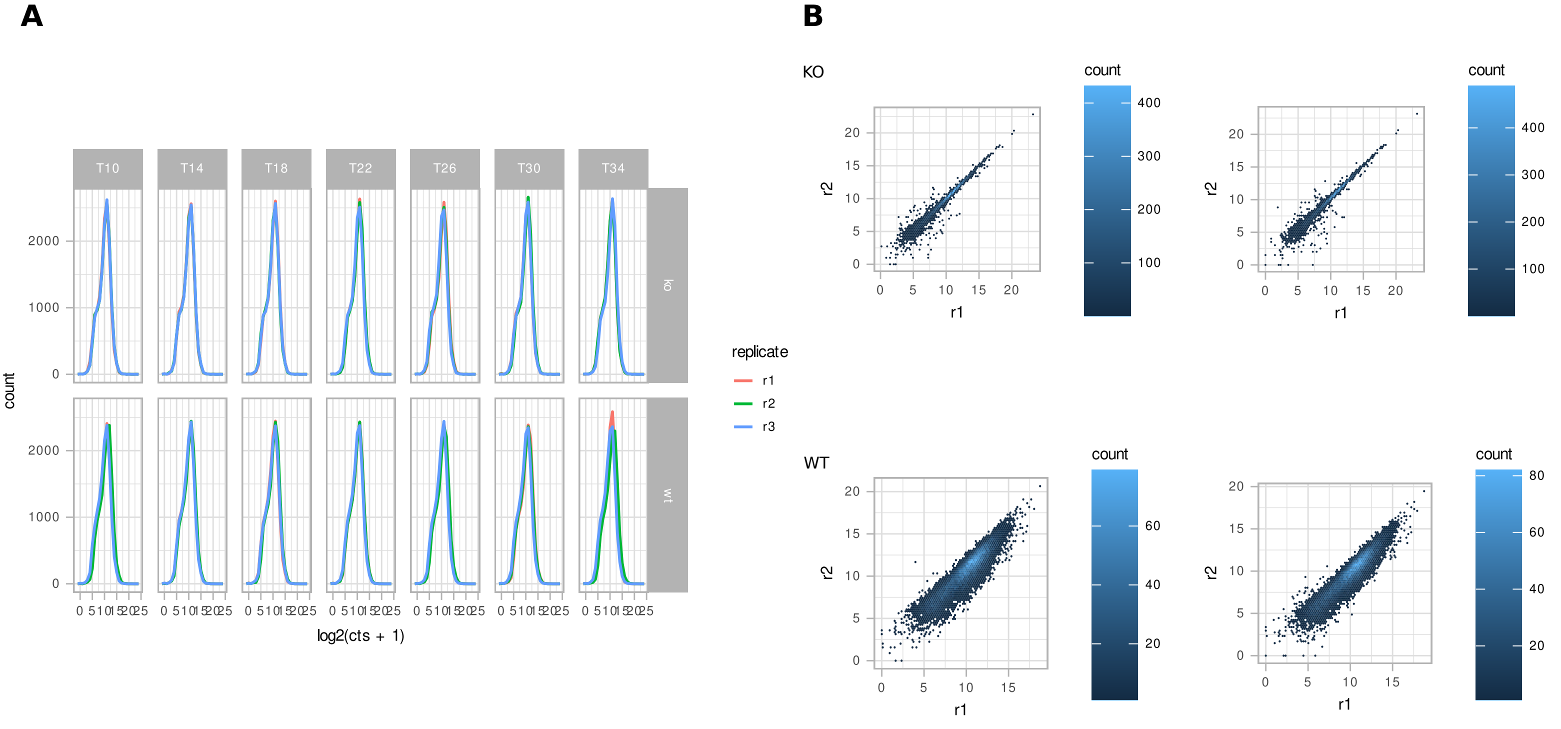


Supplemental Figure 1: Data visualization and Sample relationships Gene expression levels are estimated with featureCounts and transformed using y=log2(n+1). A. Frequence distribution of the 3 replicates r1, r2, r3 for each time-point of the two groups KO and WT. B. Correlation between the first replicate r1  and the two other replicates r2, and r3 at T34 for the two groups KO and WT
