## Supplemental Figure 2 for "Genome-wide screening of circadian and non-circadian impact of Neat1 genetic deletion"

**A.**

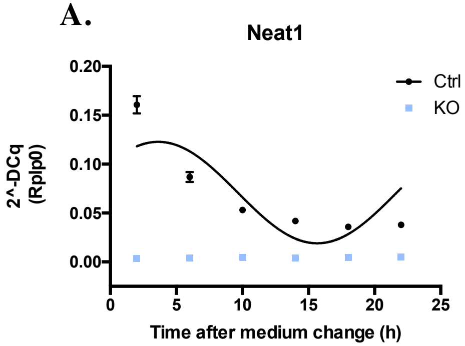


**B.**

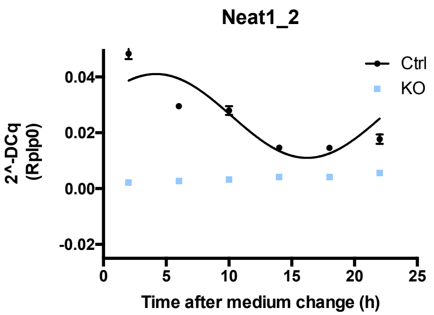


**C.**

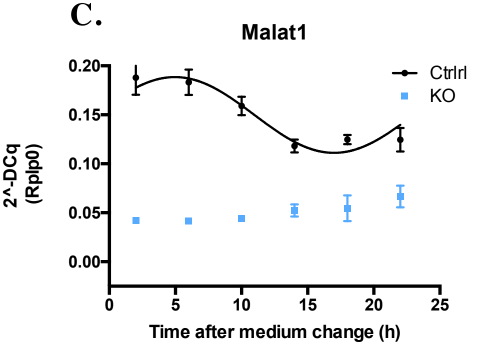


**D.**

**Cosinor fit values**

| | | Baseline | Amplitude | Phase-Shift | $R^2$ |
|---|---|---|---|---|---|
| **Neat1** | **WT** | 0,07093 | 0,05188 | 0,6253 | 0,6276 |
| **Neat1_2** | **WT** | 0,02603 | 0,01502 | 0,4769 | 0,7192 |
| **Malat1** | **WT** | 0,1498 | 0,03868 | 0,2769 | 0,6845 |

**E.**

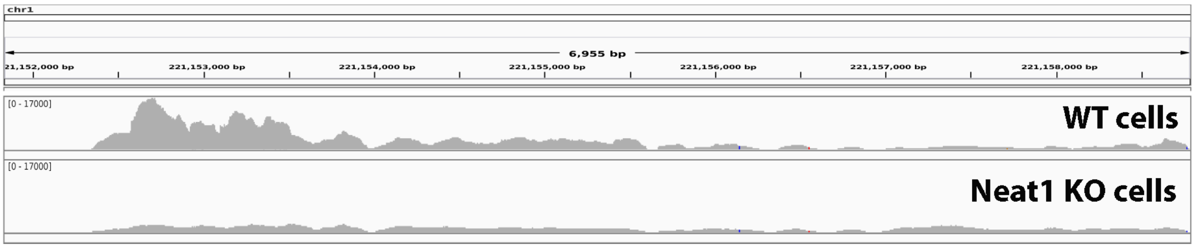


**Supplemental Figure 2: Down regulation and lost of circadian pattern of Neat1, Neat1_2 and Malat1 in Neat1 KO cells. A-C** At each time point, data are means ± SEM of three RT- qPCR measurements. In WT cells, experimental values for Neat1, Neat1_2 and Malat1 can be adequately fitted (R2>0.50) with a non-linear cosinor equation whose values are given in **D** and in which the period value is set to 24 hr (**A-C**). In Neat1 KO cells, expression levels of Neat1 and Neat1_2 is near 0 (**A, B**) and expression level of Malat1 is greatly down-regulated (**C**); values cannot be fitted by a non-linear cosinor equation in Neat1 KO cells. **E.** RNA-Seq peaks for the Malat1 locus, which is not officially annotated in the rat genome, demonstrate decreased Malat1 expression in Neat1 KO cells.
