## Supplemental Figure 3 for "Genome-wide screening of circadian and non-circadian impact of Neat1 genetic deletion"

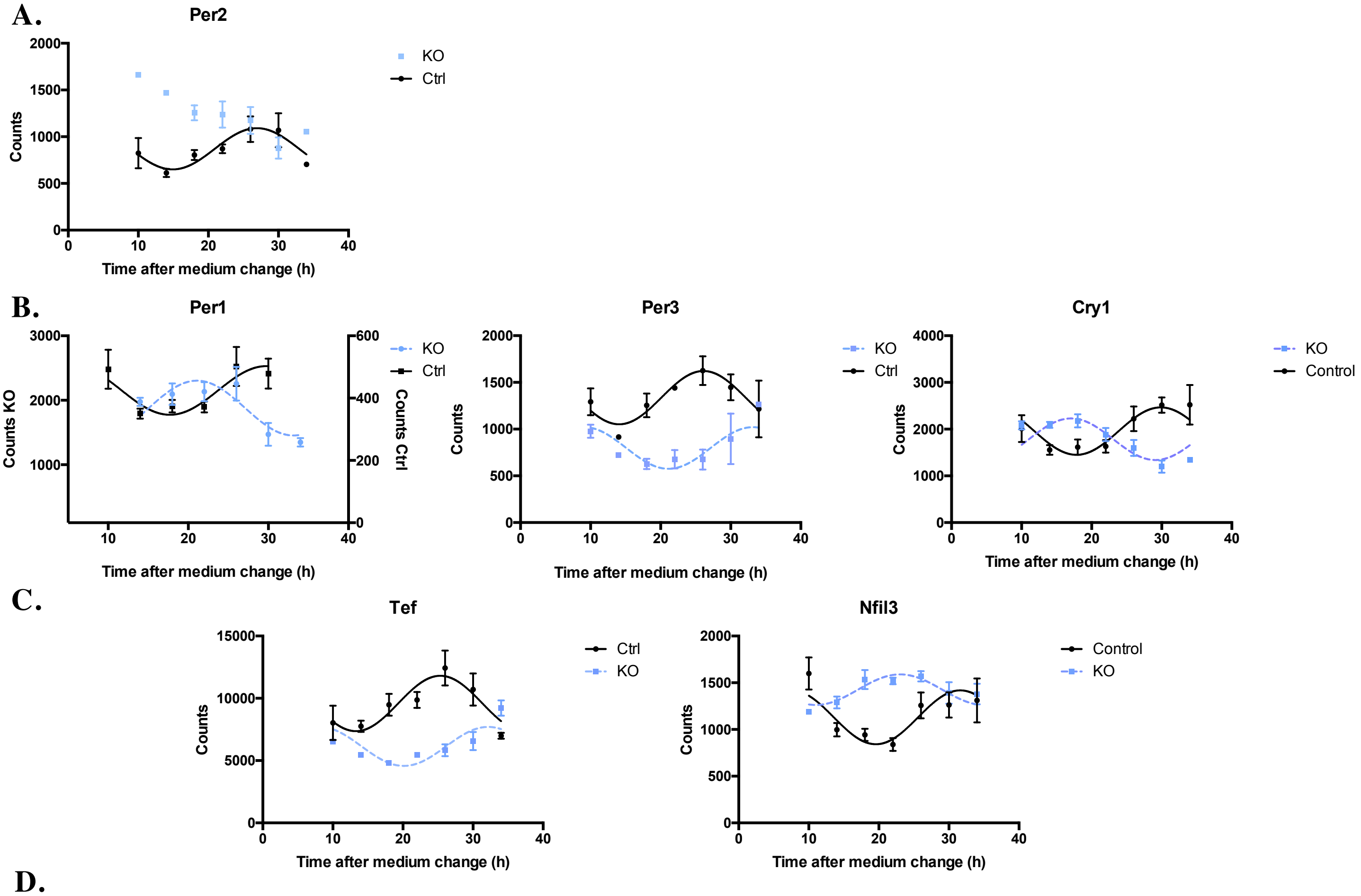


**D.**

**Cosinor fit values**

| | | Baseline | Amplitude | Phase-Shift | $R^2$ |
|---|---|---|---|---|---|
| **Per2** | **WT** | 1889 | 887 | -3.41 | 0.691 |
| **Per1** | **WT** | 1879 | -425 | -0.791 | 0.526 |
| | **KO** | 424 | 78 | 0.091 | 0.502 |
| **Per3** | **WT** | 798 | 222 | -0.798 | 0.559 |
| | **KO** | 1337 | 285 | 1.037 | 0.527 |
| **Cry1** | **WT** | 1785 | -445 | 0.257 | 0.542 |
| | **KO** | 1959 | 507 | 0.057 | 0.504 |
| **Tef** | **WT** | 9585 | 2222 | 1.242 | 0.513 |
| | **KO** | 6139 | 1565 | -0.539 | 0.536 |
| **Nfil3** | **WT** | 1426 | 163 | -17 | 0.505 |
| | **KO** | 1130 | 288 | -0.40 | 0.500 |

**Supplemental Figure 3: Impact of Neat1 genetic deletion on the circadian expression pattern of core-clock genes and direct clock-controlled genes.** At each time point, data are means ± SEM of samples from three biological replicates of RNA-Seq from WT or Neat1 KO cells. In WT cells, RNA-seq counts obtained for core-clock genes Per2, Per1, Per3 and Cry1 (**A-B**) or direct clock-controlled genes Tef and Nfil3 (**C**) can be adequately fitted ($R^2$>0.50) with a non-linear cosinor equation in which the period value was set to 24 h (cosinor fit values given in **D**). In Neat1Ko cells, while RNA-seq counts for the core-clock gene Per2 cannot be adequately fitted ($R^2$>0.50) with a non-linear cosinor equation in which the period value was set to 24 h (**A**), RNA-seq counts for the core-clock genes Per1, Per3 and Cry1 (**B**) and for the direct clock-controlled genes Tef and Nfil3 (**C**) can be adequately fitted ($R^2$>0.50) with cosinor equation (cosinor fit values given in **D**); however the circadian expression pattern of these latter genes is phase-inverted compared to their profile in WT cells (**B-C**).
