## Supplemental Figure 4 for "Genome-wide screening of circadian and non-circadian impact of Neat1 genetic deletion"

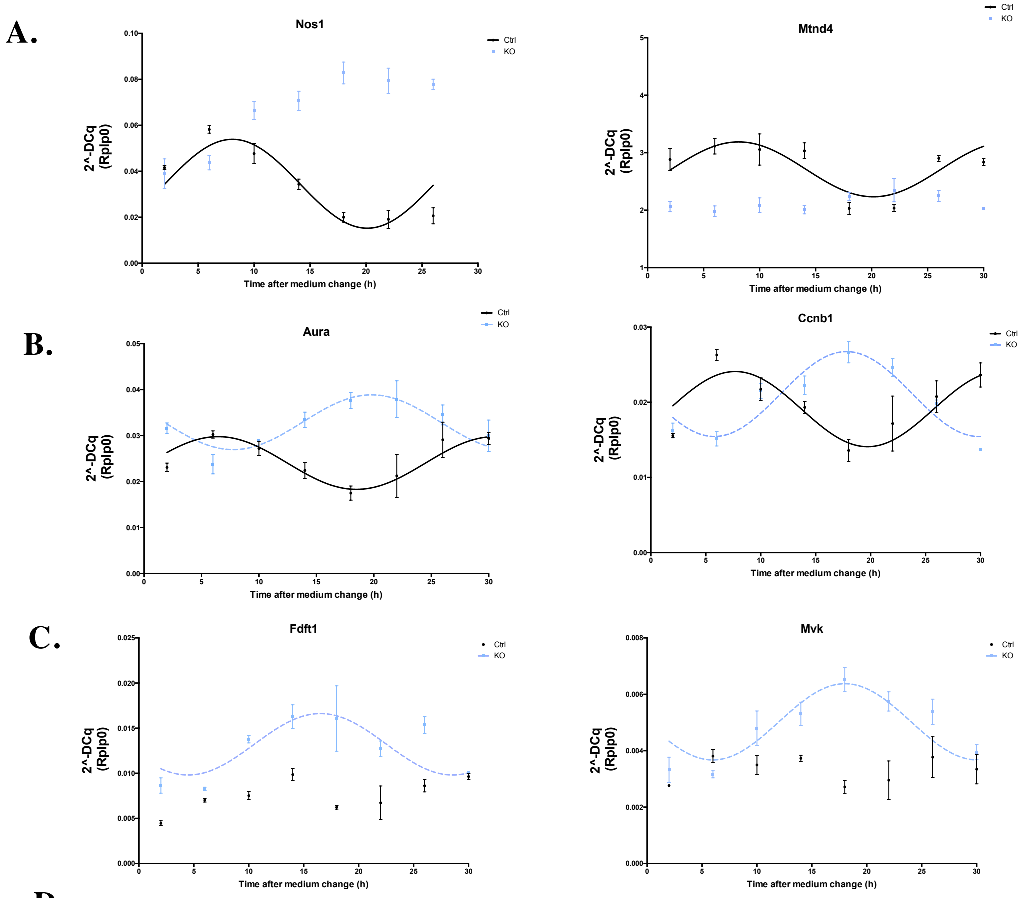


**D.**

**Cosinor fit values**

| | | Baseline | Amplitude | Phase-Shift | $R^2$ |
|---|---|---|---|---|---|
| **Nos1** | **WT** | 0.034 | 0.019 | -0.537 | 0.727 |
| **Mt-Nd4** | **WT** | 2.711 | 0.477 | -0.550 | 0.502 |
| **Aurka** | **WT** | 0.024 | 0.0057 | -0.113 | 0.525 |
| | **KO** | 0.032 | -0.0059 | -0.457 | 0.536 |
| **Ccnb1** | **WT** | 0.019 | 0.00500 | -0.434 | 0.540 |
| | **KO** | 0.021 | -0.00564 | 0.065 | 0.766 |
| **Mvk** | **KO** | 0.0050 | -0.00135 | 0.0098 | 0.570 |
| **Fdft1** | **KO** | 0.0132 | -0.00340 | 0.401 | 0.501 |

**Supplemental Figure 4: Impact of Neat1 genetic deletion on the circadian expressed genes.** At each time point, data are means ± SEM of three RT- qPCR measurements in WT or Neat1 KO cells. Loss of Neat1 impacts circadian genes in three different ways. Examples of genes that loose their rhythmic pattern after Neat1 genetic edition are given in **A**. For these genes, Nos1 and Mtnd4, experimental values from WT cells can be adequately fitted ($R^2$>0.50) with a non-linear cosinor equation in which the period value was set to 24 h (cosinor fit values given in **D**) but those from Neat1 KO cells cannot (**A**). Examples of genes that display a different circadian expression pattern in WT and Neat1KO cells are given in **B**. For these genes, Aura and Ccnb1, experimental values from WT and Neat1 KO cells can be adequately fitted ($R^2$>0.50) with a non-linear cosinor equation in which the period value was set to 24 h (cosinor fit values given in **D**); however the circadian expression pattern of these genes is phase-inverted in Neat1 KO cells compared to their profile in WT cells (**B**). Examples of genes that acquire a circadian pattern after Neat1 genetic deletion is given in **C**. For these genes, Fdft1 and Mkv, experimental values from Neat1 KO cells can be adequately fitted ($R^2$>0.50) with a non-linear cosinor equation in which the period value was set to 24 h (cosinor fit values given in **D**) but those from WT cells cannot (**C**).
