## Supplemental Table 4 for "Genome-wide screening of circadian and non-circadian impact of Neat1 genetic deletion"

**Cosinor fit values of the Fig. 2B**

|  | **Baseline** | **Amplitude** | **Phase-Shift** | **R^2^** |
| --- | --- | --- | --- | --- |
| **Alcam WT** | 13681 | 4978 | 0.9253 | 0.616 |
| **Hexim1 WT** | 3678 | 836 | -0.880 | 0.572 |
| **Cdc20 WT**  **KO** | 5137  5070 | 1270  -1074 | 0.221  -0.465 | 0.543  0.608 |
| **Nsf WT**  **KO** | 7234  9280 | 1384  -2203 | -1.044  -0.924 | 0.518  0.626 |
| **Ets1 KO** | 1016 | -156 | -0.747 | 0.662 |
| **Zfp39 KO** | 355 | 140 | 1.054 | 0.715 |

**Supplemental Table 4: Cosinor analysis of the rhythmic expression pattern of different genes differentially affected by the Neat1 deletion.** Mean experimental values ± SEM of three RT- qPCR measurements in WT or Neat1 KO cells are fitted with a non-linear sine wave equation (Y = Baseline + Amplitude * sin (Frequency*X + Phase-shift) in which the period value (2pi/Frequency) is constrained to 24h. Experimental values are considered well fitted by cosinor regression when the R squared was higher than 0.50.
